# Role of *AtCPK5* and *AtCPK6* in the regulation of the plant immune response triggered by rhamnolipids in Arabidopsis

**DOI:** 10.1101/2025.10.22.683368

**Authors:** Juliette Stanek, Olivier Fernandez, Marie Boudsocq, Dina Aggad, Sandra Villaume, Laetitia Parent, Sandrine Dhondt Cordelier, Jérôme Crouzet, Stéphan Dorey, Sylvain Cordelier

## Abstract

Rhamnolipids (RLs) are a class of glycolipids naturally synthesized by bacteria potentially used in the biocontrol of plant pathogenic microorganisms in agriculture. While RLs trigger an immune response in plants, little is known about the signaling mechanisms involved after RL perception. Calcium Dependent Protein Kinases (CDPKs/CPKs) are a large family of kinases involved in various functions in plants including the regulation of the resistance response to phytopathogens. Here, we investigated the role of *AtCPK5* and *AtCPK6* in RL-triggered immunity in *Arabidopsis thaliana* (hereafter referred to as Arabidopsis). Gene expression analysis showed that RLs up-regulate the expression of both genes in Arabidopsis leaves. Using a functional approach, we demonstrated that signaling events exemplified by reactive oxygen species (ROS) production and defense gene expression including *AtWRKY46*, *AtFRK1* and *AtPR1* are increased in *cpk5/6* mutant plants compared to wild-type (WT) plants. Moreover, *cpk5* mutants displayed intermediary responses while *cpk6* mutation alone did not change the immune profile activated by RLs, except for *AtFRK1*. Those results suggest that *AtCPK5* and *AtCPK6* regulate RL-triggered defenses, with *AtCPK5* likely playing a more important role. Yet, the *cpk5/6* mutations did not affect RL-triggered electrolyte leakage nor enhanced RL-mediated resistance to the bacterial pathogen *Pseudomonas syringae* pv tomato DC3000. Our results therefore show that additional signaling components regulate *AtCPK5* and *AtCPK6*-mediated plant immune responses triggered by RLs.

## Introduction

Under natural conditions, plants are subject to various biotic stresses such as pathogen attacks (Zhang *et al*., 2022; Du *et al*., 2024). In order to overcome these aggressions, they are able to activate defense mechanisms, which are largely based on the activation of an effective immune response (Zia *et al*., 2021; Ngou *et al*., 2022*a*; Jones *et al*., 2024). As a first key step of this immune response, plant cells detect pathogens through the recognition of Invasion Patterns (Ips ; Schellenberger *et al*., 2019) by Pattern Recognition Receptors (PRRs) activating the so-called Pattern-Triggered Immunity (PTI ; Ngou *et al*., 2024). Following this recognition phase, different signaling events occur, including the production of Reactive Oxygen Species (ROS), phosphorylation/dephosphorylation cascades and ion fluxes (Cordelier *et al*., 2022; Ngou *et al*., 2022*a*,*b*). The influx of calcium ions (Ca²⁺) represents one of the key signaling events in plant immune responses. Transient cytosolic [Ca²⁺] elevations function as secondary messengers translating pathogen perception into appropriate defense responses (Verma *et al*., 2022; Mariyam *et al*., 2023). When a given stress condition arises, a significant release of Ca2+ ions occurs into the cytosol. Additionally, a specific Ca2+ signature can be associated to specific biotic and abiotic stresses (Boudsocq and Sheen, 2010; Liese *et al*., 2024; Wdowiak *et al*., 2024). These ions are detected by various Ca2+ sensors such as calmodulins (CaMs), calcium and calmodulin-dependent protein kinases (CCaMKs), calcineurin B-like proteins (CBLs) and calcium-dependent protein kinases (CDPKs/ CPKs) (Boudsocq and Sheen, 2010; Aldon *et al*., 2018; Kudla *et al*., 2018). Ca2+ modifies the structural conformation and/or enzymatic activity of these Ca2+ sensors, which activates target proteins to transfer signals to downstream pathways.

CPKs are involved in a wide range of functions, including plant development (roots and shoots) and plant responses to biotic and abiotic stresses (Dekomah *et al*., 2022). In Arabidopsis, there are 34 different CPKs (Harmon *et al*., 2001). CPKs share a four-part structure including a variable N-terminal domain, a Serine/Threonine protein kinase domain, an auto-inhibitory junction domain and a C-terminal calmodulin-like domain (EF-hand domain) (Dekomah *et al*., 2022). Their activation is triggered by cytosolic Ca²⁺ to their EF-hand domains (Liese and Romeis, 2013; Kiselev and Dubrovina, 2025). Based on sequence similarity, particularly within the catalytic kinase domain and the regulatory calmodulin-like domain, CPKs are phylogenetically classified into four subgroups (I-IV). This classification may reflect functional diversification and evolutionary adaptation to different signaling pathways (Wang *et al*., 2016; Mu *et al*., 2024). CPKs functions can be quite distinct from one sub-group to another. To date, 10 CPKs have been shown to be involved in biotic stress responses in Arabidopsis (Bredow and Monaghan, 2019). For example, *At*CPK1 acts as a positive regulator of resistance against pathogens with different lifestyles, such as *Botrytis cinerea or Pseudomonas syringae pv tomato* DC3000 (hereafter referred to as *Pst* DC3000; Coca and San Segundo, 2010). Both *At*CPK4 and *At*CPK11, are known to act as positive regulators of pathogen responses during Effector-Triggered Immunity (ETI) and PTI by regulating defense gene expression and *At*RBOHD phosphorylation (Gao *et al*., 2014; Yip Delormel and Boudsocq, 2019). Several authors have emphasized the redundant action of *At*CPK5 and *At*CPK6 under biotic stress conditions (Boudsocq *et al*., 2010). They are involved in phosphorylation of transcription factors like WRKYs and the induction of antimicrobial metabolites such as camalexin (Gao *et al*., 2013; Zhou *et al*., 2020*a*). The loss-of-function Arabidopsis double mutant *cpk5/6* showed reduced resistance to pathogens like *B. cinerea* (Gravino *et al*., 2015; Yang *et al*., 2020)*, Pst DC3000, Pst avrRpm1* and *Pst avrRpt2* (Boudsocq *et al.,* 2010; Gao *et al.,* 2013), highlighting their redundant and essential roles in PTI and ETI. They are also involved in Programmed Cell Death (PCD) and ethylene biosynthesis in response to *B. cinerea* (Gravino *et al*., 2015; Yang *et al*., 2020; Zhou *et al*., 2020*a*).

Over the last decade, amphiphilic IPs, such as glycolipids and lipopeptides, have been investigated as novel natural bio-based molecules for the biocontrol of plant diseases (Crouzet *et al*., 2020; Furlan *et al*., 2020). Rhamnolipids (RLs) are natural, highly biodegradable molecules that can induce disease resistance to phytopathogens in various plant species (Crouzet *et al*., 2020). Natural RLs activate an immune response in grapevine, rapeseed and in Arabidopsis (Varnier *et al*., 2009; Sanchez *et al*., 2012; Monnier *et al*., 2018). In Arabidopsis, the lipid tail of RLs (HAAs, for (R)-3-hydroxyalkanoates) is perceived by the bulb-type lectin receptor kinase LIPOOLIGOSACCHARIDE-SPECIFIC REDUCED ELICITATION/S-DOMAIN-1-29 (LORE/SD1-29), which also mediates medium-chain 3-hydroxy fatty acids (mc-3-OH-FAs) sensing. On one hand, HAAs and mc-3-OH-FAs trigger an immune response exemplified by an early ROS production (Kutschera *et al*., 2019; Schellenberger *et al*., 2021). On the other hand, RLs elicit LORE-independent defense responses (Schellenberger *et al*., 2021). This RL-induced immune response is characterized by the up-regulation of classical defense genes (Varnier *et al*., 2009; Sanchez *et al*., 2012) and in a non-canonical ROS signature displayed by a late and sustained *At*RBOHD-dependent ROS production (Schellenberger *et al*., 2019). The receptor-like cytoplasmic kinase *At*BIK1, which often regulates the activation of *At*RBOHD (Wang *et al*., 2020), is not involved in RL-induced response (Schellenberger *et al*., 2019).

Since *At*RBOHD can also be activated by phosphorylation from subgroup I CPKs, especially *At*CPK5 and *At*CPK6 (Dubiella *et al*., 2013; Kadota *et al*., 2014), we investigated in this work the role of these two CPKs, in the plant immune response triggered by RLs. First, we demonstrated that RLs trigger the expression of *AtCPK5* and *AtCPK6* in Arabidopsis leaves. Using a functional approach, we found that a synergistic effect of *At*CPK5 and *At*CPK6 negatively regulates ROS production and defense gene expression after RL perception. We also showed that those genes are not directly involved in the RL-triggered local resistance to the hemibiotrophic bacterial pathogen *Pst* DC3000. Therefore, our results highlight the involvement of additional proteins in CPK-mediated immune response activated by these amphipathic elicitors.

## Material and Methods

### Plant material and molecules

Arabidopsis ecotype Col-0 was used as wild-type (WT) parent for all experiments. Seeds from *cpk5* (sail_657_C06), *cpk6* (salk_025460) *and cpk5/6* (sail_657_C06, salk_025460) Arabidopsis homozygous mutants were previously reported (Boudsocq *et al*., 2010). Seeds from *sd1-29* (*lore-5*), Col-0AEQ, and *lore-5*AEQ Arabidopsis homozygous mutants were provided by S. Ranf (Ranf *et al*., 2015, Schellenberger *et al.,* 2021). All Arabidopsis mutants are in the Col-0 background. Plants were grown on Presstopf Tray soil (Gramoflor, Vechta Niedersachsen, Germany) in growth chambers at 20°C, under 12-h light/12-h dark regime and 60% relative humidity. RLs (AGAE® Company, USA) were used at 0,6 mg/mL in water (Sanchez *et al*., 2012; Schellenberger *et al*., 2021) for all experiments.

### Conductivity assay

Conductivity assays were carried out on 5-to 6-week-old Arabidopsis plants cultured on soil in growth chamber. Four leaf discs of 6mm diameter were incubated in distilled water for 2 hours. Two discs were placed in a well of a 12-well plate (Falcon®) containing fresh distilled water with the RLs and two in water for control. Conductivity measurements (three replicates for each treatment) were then conducted 24 hours after treatment using a B-771 LaquaTwin (Horiba) conductivity meter.

### Extra-cellular ROS production and Ca2+ influx analysis

ROS assays were carried out on 5-to 6-week-old Arabidopsis plants cultured on soil in growth chamber. Leaf discs of 4mm diameter were cut and placed in 150 μL distilled water overnight in a 96-well plate (Falcon®). The following processes were made as detailed by (Smith and Heese, 2014). Luminescence (Relative Light Units, RLU) was measured every 5 min during 12 hours with a TECAN SPARK Multimode Microplate Reader. Control was realized on leaf discs of WT plants with water. Ca2+ influx analysis were performed using Col-0AEQ and *lore-5*AEQ mutant as detailed by (Ranf *et al*., 2015). Luminescence measurements were performed following the same procedure with a TECAN SPARK Multimode Microplate Reader (TECAN®).

### Gene expression analysis

In our assays, leaf discs of 6mm diameter of 5-to 6-week-old Arabidopsis plants were cut and placed in 2 mL distilled water for 2 hours in a 24-well plate (Falcon®). They were collected at 0 hours, 9 hours and 24 hours after RL treatment and crushed with liquid nitrogen. 50 mg of each sample were placed in 2 mL cold safe-lock Eppendorf. 1 mL of QIAzol Lysis Reagent (QIAGEN®) were dropped in each tube and were vortexed. After 5 minutes of incubation, chloroform/IAA (24:1) was added. All the samples were vigorously shaken and centrifuged for 15 minutes at 10,000 g. The aqueous phase is collected and precipitated with isopropanol then centrifuged at 10,000 g. Samples were washed with 70% ethanol and pellets were solubilized in purified water DNAse/RNAse free. cDNAs were obtained following a Reverse Transcriptase protocol (EuroBlueTaq kit).

Quantitative RT-PCR was carried out on three independent biological replicates for each sample, as well as two technical replicates for each reaction. Quantitative RT-PCRs were performed using qPCRBIO SyGreen Blue Mix Lo-Rox (Eurobio Scientific, France, Les Ulis) in white 384-well plates (Sorenson, USA, Salt Lake City) and on a CFX Opus 384 instrument (Bio-Rad, USA, Hercules). Expression levels of the selected genes were normalized to *AtActin7* (GenBank NM_121018.4 ; Cao *et al*., 2008), *AtActin2* (GenBank NM_112764.4 ; Yuan *et al*., 2025) and *AtTubulin4* (GenBank NC_003074.8 ; Körber *et al*., 2016) expression as reference genes. The primers used in this study are listed in supplemental table 1 and some were previously reported (Boudsocq *et al*., 2010).

### Pst DC3000 culture and disease-resistance assays

*P. syringae pv. tomato* strain DC3000 (*Pst* DC3000) was grown at 28 °C under stirring in King’s B (KB) liquid medium supplemented with antibiotic: 50 μg/mL rifampicin. For protection assays, Arabidopsis plants were grown individually for 4 weeks in soil. For each experiment, six pots per condition were used (n = 6). The following processes were made as detailed in (Schellenberger *et al*., 2021). Two days before infection, plants were sprayed with RLs or water as control and were placed in high humidity atmosphere. Plants were then infiltrated with bacterial suspension at the concentration of 10^7^ CFU/mL (in 10 mM MgCl2) using a needleless syringe. Bacterial quantification *in planta* (colony forming units; CFU) was performed 3 days post infection (dpi). To this end, all plant leaves from the same pot were harvested, weighed, and crushed in a mortar with 10 mL of 10 mM MgCl2, and serial dilutions were performed. For each dilution, 10 μL were dropped on KB plate supplemented with appropriate antibiotics. CFU were counted after 2 days of incubation at 28 °C. The number of bacteria per milligram of plant fresh mass was obtained with the following formula:

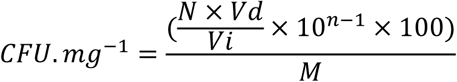

with N equal to CFU number, Vi the volume depot on plate, Vd the total volume, n the dilution number, and M the plant fresh mass.

### Statistical analysis

All data were subjected to statistical analysis using different tests with the Rstudio software. “Vegan” package was used for the PCoA analysis and “stats” package was used for the other analyses. GraphPad software was used for graphical representation of data.

## Results

### RLs trigger Ca²⁺ influx in Arabidopsis leaves

We first investigated Ca²⁺ influx following RL challenge, as it is an essential component for CPK activation (Xu *et al*., 2024). RLs are known to induce a strong, late and sustained production of ROS in Arabidopsis leaves from 3 to 12 hours post treatment (hpt ; Schellenberger *et al*., 2021). Similarly, we observed a strong and late influx of Ca²⁺ in Col-0Aeq Arabidopsis plant after challenge with RLs, starting at 3hpt and decreasing at 10hpt (fig1A). A Ca²⁺ influx was also observed in *lore-5*Aeq plants that does not sense 3-OH-FA and HAA precursors, which could be found in trace amount in RL solution (fig1B ; Schellenberger *et al*., 2021). These results clearly demonstrate that RLs are inducing Ca²⁺ influx in Arabidopsis.

### RLs trigger up-regulation of *AtCPK5* and *AtCPK6* genes expression

*AtCPK5* and *AtCPK6* gene expression was examined in WT and in *lore-5* plants after treatment with RLs (fig. 2). RT-qPCR results show a significant increase in transcripts of *AtCPK5* and *AtCPK6* is both WT and mutant plants 24h after RL challenge with no significant differences in both plant backgrounds (fig. 2A & B). The relative expression of *AtCPK5* at 9 hpt and 24 hpt in WT, *cpk5*, *cpk6* and *cpk5/6* plants revealed that the gene was induced to a similar extent in WT and in *cpk6* mutants following RL treatment (fig. 3A). As expected, no expression of *AtCPK5* was observed in the *cpk5* and *cpk5/6* mutant plants. Similarly, there was no significant difference in *AtCPK6* gene expression levels between WT plants and *cpk5* mutant plants both treated with RLs (fig. 3B). No *AtCPK6* expression was detected in both *cpk6* and *cpk5/6* plants.

**Figure 1.**
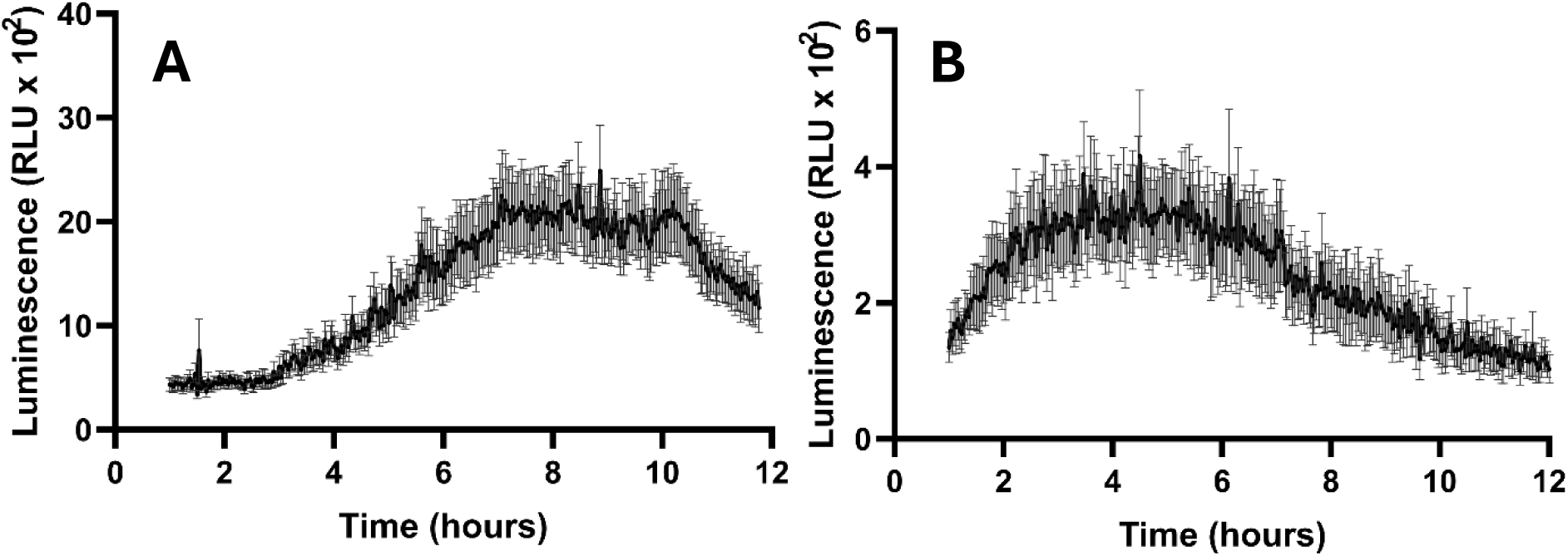
RLs induce late cytoplasmic [Ca2+] production in Arabidopsis leaf. [Ca2+]cyt of WT-Aeq (**A**) and *lore-5*-Aeq (**B**) was quantified in Arabidopsis leaves after RL treatment. RLU amounts were analysed over 12 hours. Data are presented as mean ± SEM (n=7, experiments were performed two times with similar results).

**Figure 2.**
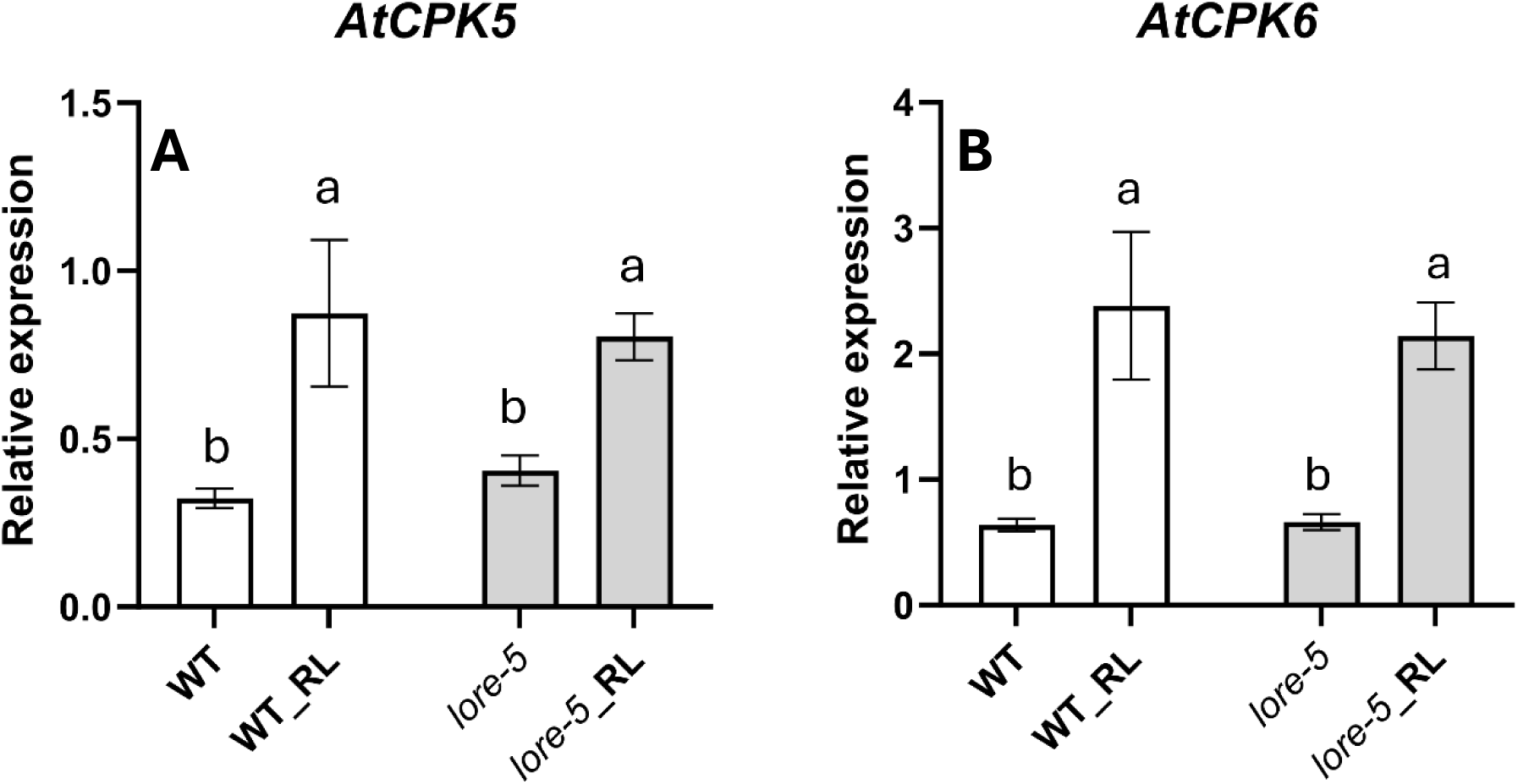
RLs induce *AtCPK5* and *AtCPK6* gene expression in Arabidopsis leaf. RL-induced *AtCPK5* (**A**) and *AtCPK6* (**B**) gene expression was studied by RT-qPCR in WT plants and *lore* mutants. Leaf discs were treated with RLs (0.6 mg/mL) and analysed at 24 hpt. Data are presented as mean ± SEM (n=6, three independent experiments). Expression data were normalized with control (non treated WT at 0hpt) and compared with *AtActin2* and *AtTubulin4* as reference genes. Letters represent results of Kruskal-Wallis followed by Wilcoxon pairwise test by time, with *P* > 0.05 (same letters) or *P* ≤ 0.05 (different letters).

**Figure 3.**
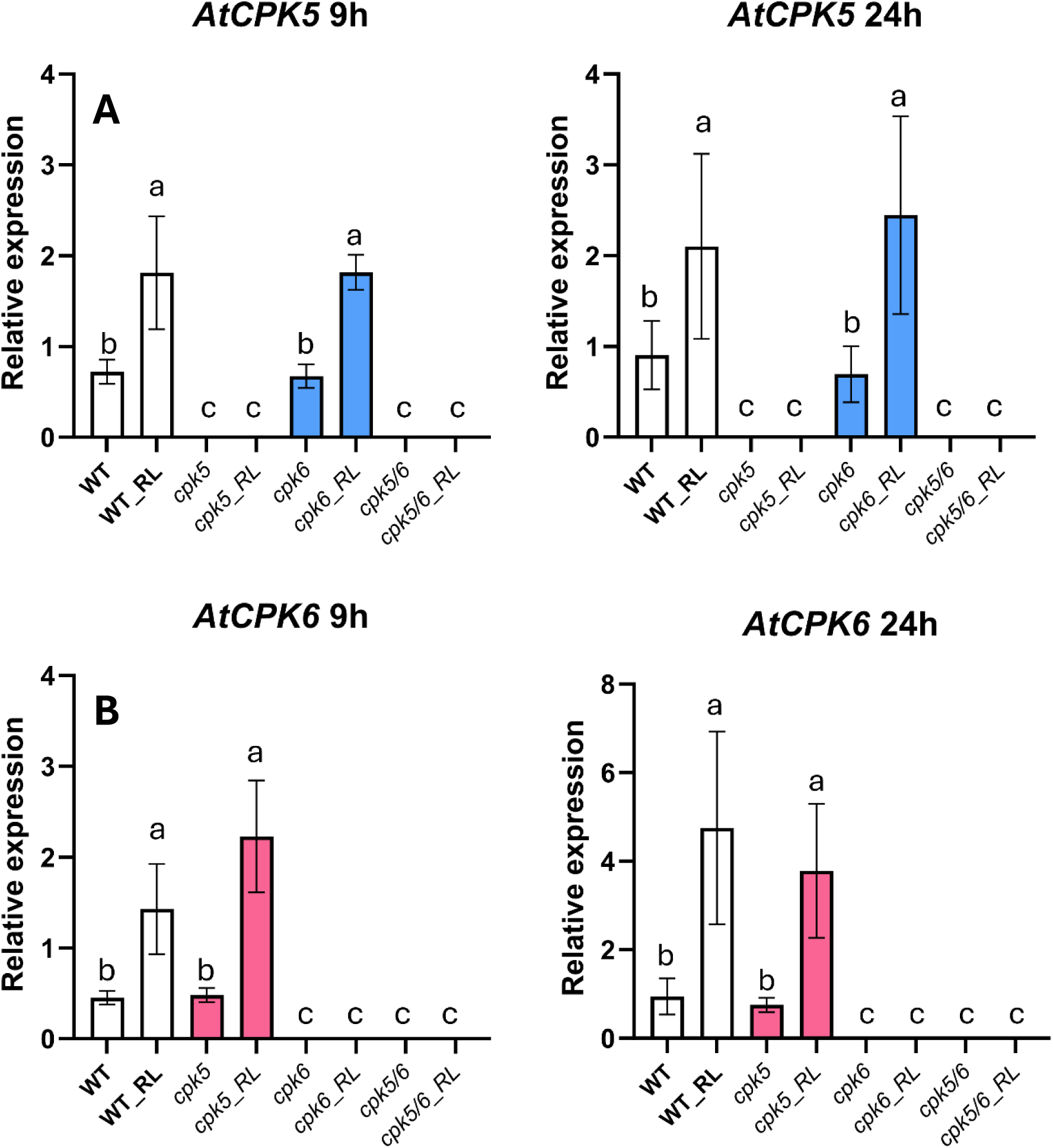
RLs independently induce *AtCPK5* and *AtCPK6* gene expression. *AtCPK5* (**A**) and *AtCPK6* (**B**) gene expression following RL treatment was analysed by RT-qPCR in WT plants and *cpk5*, *cpk6* and *cpk5/6* mutants. Leaf discs were treated with RL (0.6 mg/mL) and analysed at 0hpt, 9 hpt and 24 hpt (0hpt data are not presented here). Expression data were normalized with *AtActin7*, *AtActin2* and *AtTubulin4* as reference genes. Data are presented as mean ± SEM (n=6, three independent experiments). Letters represent results of Kruskal-Wallis followed by Wilcoxon pairwise test by time, with *P* > 0.05 (same letters) or *P* ≤ 0.05 (different letters).

### Extracellular ROS production is increased in *cpk5/6* background following RL perception

We compared ROS production levels after RL treatment from 1 to 12 hpt (corresponding to the typical RL-triggered ROS signature; Schellenberger *et al*., 2021) in WT and mutant backgrounds (fig. 4A). ROS production in *cpk6* leaves was similar to WT leaves, while a little higher in *cpk5*. However, the production of extracellular ROS was strongly enhanced in the *cpk5/6* double mutant leaves (fig. 4A). To complete our statistical analysis of the ROS production, we performed a PCoA analysis, based on the calculation of the Frechet distance between each measurement for each condition at each time point in the series. The following PERMANOVA analysis on the resulting intergroup distance allowed us to conclude that only the *cpk5/6* double mutant exhibited significantly elevated levels compared to WT (fig. 4B). This analysis indicates that both *At*CPK5 and *At*CPK6 negatively regulate the ROS burst induced by RLs, with *At*CPK5 likely playing a major role.

**Figure 4.**
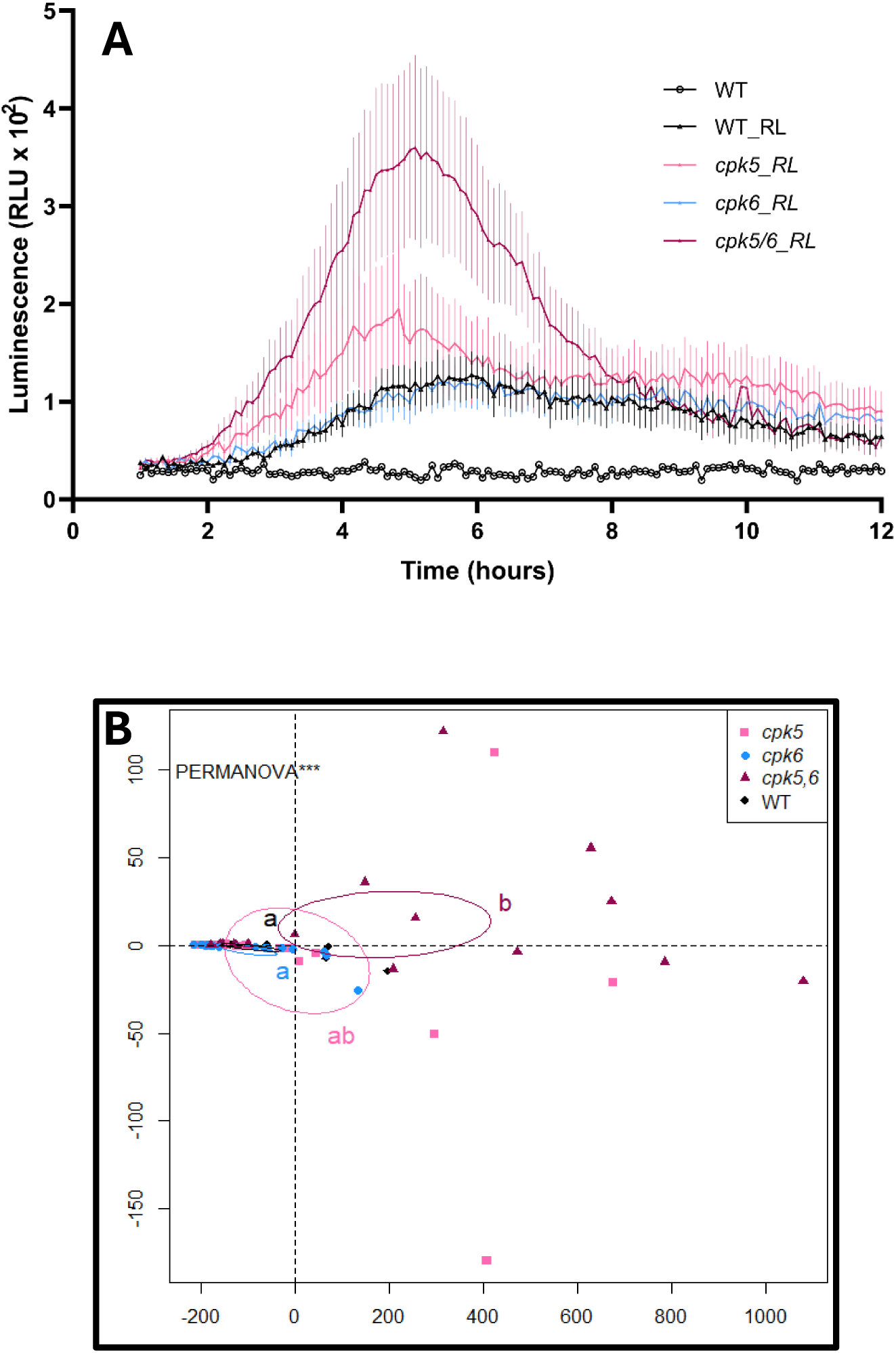
*At*CPK5 and *At*CPK6 are involved in ROS production after RL treatment in Arabidospis leaf. (**A**) Extracellular ROS production after treatment of WT, *cpk5*, *cpk6* and *cpk5/6* mutants leaf discs with 0.6 mg/mL RLs or purified water as control. ROS production was analysed from 1 hpt to 12 hpt. (***B***) Principal Coordinate Analysis (PCoA) plot deduced from Frechet distances between temporal measurements of ROS production during the second peak following RLS mix treatment. The PERMANOVA test (and its pairwise version using FDR correction) was used for statistical analysis. Different letters indicate statistically different groups (p <0.01). Ellipses display confidence intervals (95%). (*A*–*B*) Data are mean ± SEM (*n* = 18, three independent experiments).

### *At*CPK5 and *At*CPK6 are involved in regulation of defense genes activated by RLs

To determine whether *At*CPK5 and *At*CPK6 would potentially be involved in the regulation of defense genes following RL challenge, we analysed the expression of *AtWRKY46* (NM_130204.3) and *AtFRK1* (Flg22-Induced Receptor-Like Kinase 1, NM_127476.2) as early defense markers known to be involved in classical PTI (He *et al*., 2006; Biniaz *et al*., 2022) and *AtPR1* (Pathogenesis-Related 1, NM_127025.3) as a late defense marker. *AtPR1* is a classical defense-related gene that is widely used as a molecular marker for the salicylic acid signaling pathway, and resistance to biotrophic pathogens (Laird *et al*., 2004; Pečenková *et al*., 2022; Hao *et al*., 2024; Kovács *et al*., 2025). *AtWRKY46* gene was slightly upregulated at 9 hpt in RL-treated WT plants (fig. 5A). Compared to WT, a 3 and 6-fold increase in *AtWRKY46* expression was respectively observed in *cpk5* and *cpk5/6* mutants following RL challenge. No difference was found in *AtWRKY46* induction between WT and *cpk6* mutant (fig. 5A). However, *At*CPK6 also appears to be involved, since *AtWRKY46* is more highly expressed in the *cpk5/6* double mutant than in the *cpk5* single mutant. Following RL treatment, the expression pattern of *AtFRK1* gene was similarly increased in *cpk5, cpk6* and *cpk5/6* mutants (fig. 5B). *AtPR1* expression was up-regulated in the *cpk5* and *cpk5/6* mutants but not in *cpk6* mutant after RL challenge (fig. 5C). However, as with the *AtWRKY46* gene, *At*CPK6 also appears to be involved, since *AtPR1* is more highly expressed in *cpk5/6* than in *cpk5*.

**Figure 5.**
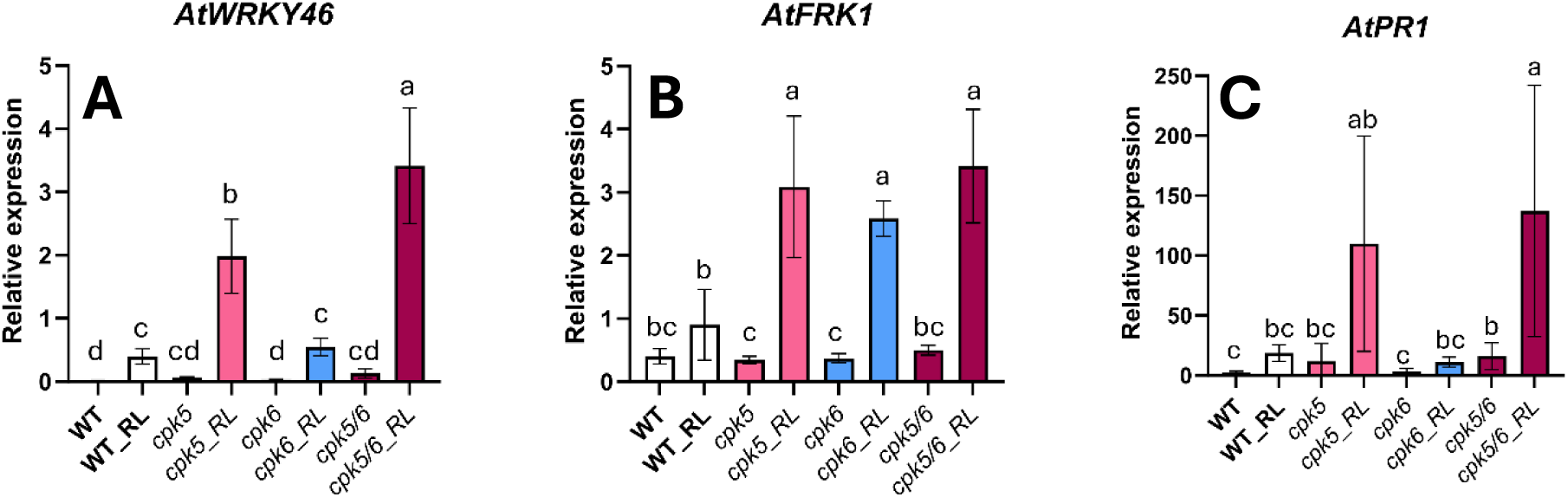
*At*CPK5 and *At*CPK6 are involved in early and late defense gene expression after RL treatment. *AtWRKY46* (**A**), *AtFRK1* (**B**) and *AtPR1* (**C**) gene expression was studied by RT-qPCR in WT plants and *cpk5*, *cpk6* and *cpk5/6* mutants. Leaf discs were treated with RLs (0.6 mg/mL) and analysed at 9 hpt for early defense gene expression (*AtWRKY46* and *AtFRK1*) and at 24 hpt for late defense gene expression (*AtPR1*). Data are presented as mean ± SEM (n=6, three independent experiments). Expression data were normalized with *AtActin7*, *AtActin2* and *AtTubulin4* as reference genes. Letters represent results of Kruskal-Wallis followed by Wilcoxon pairwise test by time, with *P* > 0.05 (same letters) or *P* ≤ 0.05 (different letters).

### *AtCPK5* and *AtCPK6* inactivation does not affect RL-triggered electrolyte leakage

Electrolyte leakage is a typical marker of RL-triggered immunity (Schellenberger *et al*., 2021). As expected, when treated with RLs, WT plants displayed a strong increase of electrolyte leakage at 24 hpt (fig. 6). This response was conserved in *cpk5*, *cpk6* and *cpk5/6* mutant plants, which showed no significant differences compared to WT plants.

**Figure 6.**
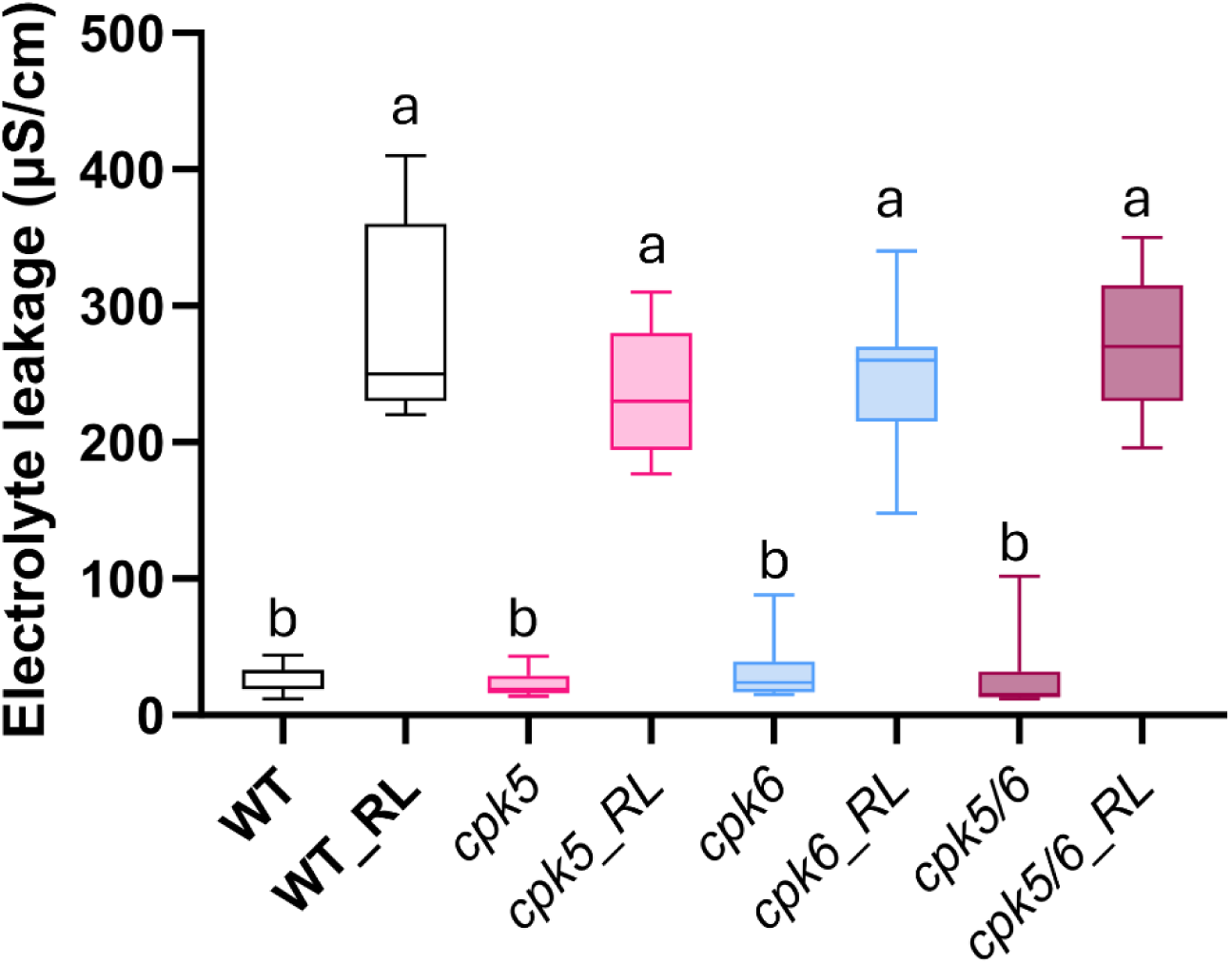
*At*CPK5 and *At*CPK6 are not involved in RL-induced electrolyte leakage on Arabidopsis leaf. Electrolyte leakage was measured on WT, *cpk5*, *cpk5* and *cpk5/6* Arabidopsis leaf discs 24 h post treatment by RLs (0.6 mg/mL) or purified water (control). Data are mean ± SEM (n=9, three independent experiments). Letters represent results of Kruskal-Wallis followed by Wilcoxon pairwise test by time, with *P* > 0.05 (same letters) or *P* ≤ 0.05 (different letters).

### *At*CPK5 and *At*CPK6 are not involved in Arabidopsis RL-triggered resistance to the hemibiotrophic pathogen *Pst* DC3000

*Pst* DC3000 is a well-studied model pathogen and is classified as a hemibiotrophic pathogen that initially feeds on living plant tissues and later causes the death of plant cells (Yang *et al*., 2023; Blomberg *et al*., 2024; Liu *et al*., 2024). To study RL-induced resistance to *Pst* DC3000, Arabidopsis plants were pre-treated with water control or RLs 2 days before bacterial infection. In control conditions, the *cpk5* and *cpk6* single mutants display a sensitivity to *Pst* DC3000 comparable to the WT while the *cpk5/6* double mutant was hypersensitive (fig 7), as previously reported (Boudsocq *et al*., 2010). Upon RL treatment, all plant backgrounds, including the three *cpk* mutants, displayed the same level of protection induced by RL treatment against *Pst* DC3000, indicating that *At*CPK5 and *At*CPK6 are not essential for RL-triggered resistance (fig. 7).

**Figure 7.**
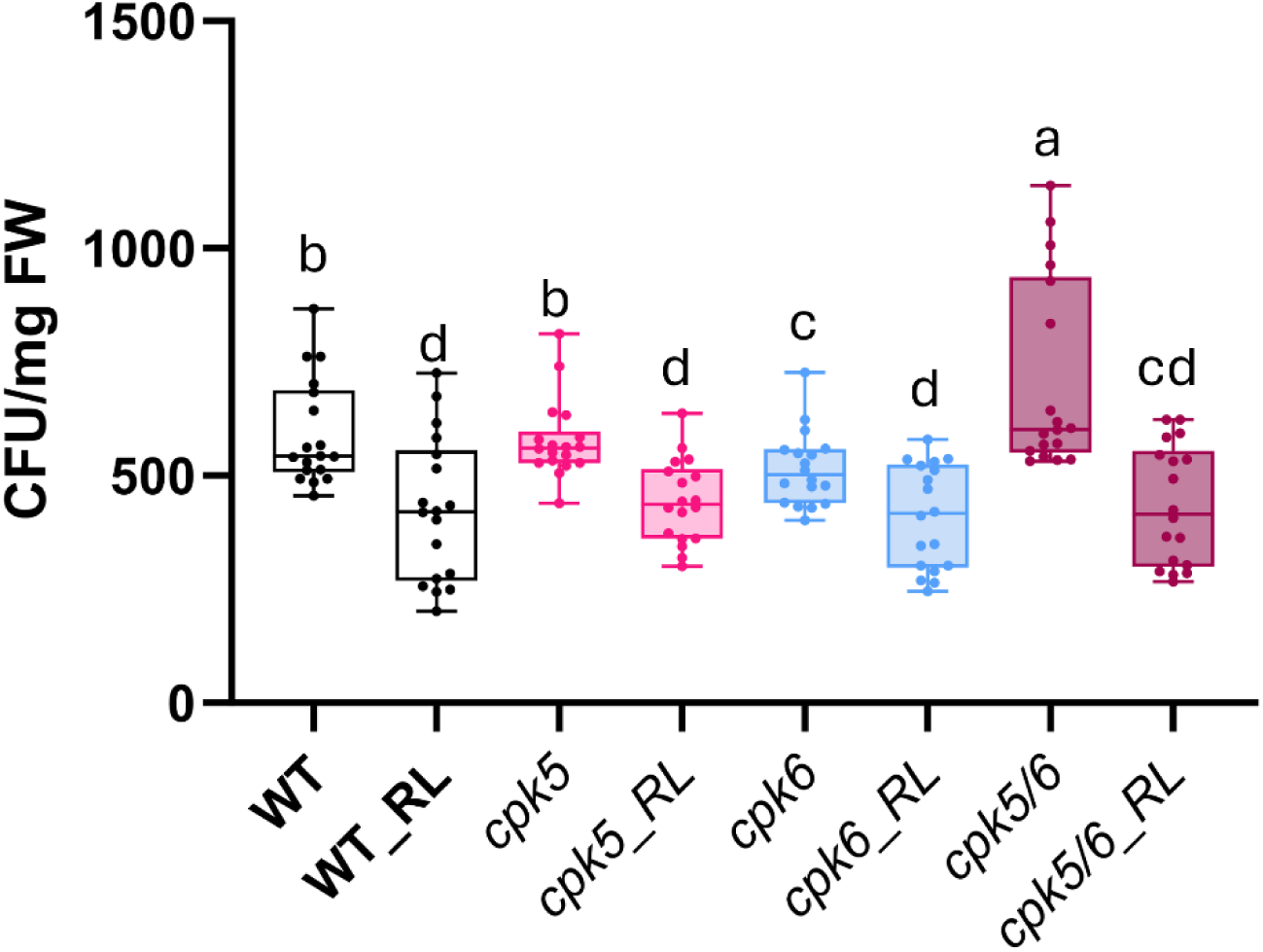
*At*CPK5 and *At*CPK6 are not involved in RL-induced *Pst* protection on Arabidopsis leaf. WT, *cpk5, cpk6* and *cpk5/6* Arabidopsis leaves were treated with RLs (0.6 mg/mL) or purified water (control) 48 h before infection. *Pst* colonies were counted at 3 dpi. Data are individual data ± SEM (*n*=18, three independent experiments). Letters represent results of Kruskal-Wallis followed by Wilcoxon pairwise test by time, with *P* > 0.05 (same letters) or *P* ≤ 0.05 (different letters).

## Discussion

In this study, we demonstrated that RLs activate Ca2+ influx in Arabidopsis. This is the first time that RLs have been shown to induce a sustained and late Ca2+ signature in Arabidopsis. This signature is similar to analyses of extracellular ROS production following RL challenge (Schellenberger *et al*., 2021). Interestingly, previous studies have suggested that these two types of signatures may be interconnected (Ravi *et al*., 2023; Weralupitiya *et al*., 2024). It has been shown that Ca²⁺ is important at different levels of plant immunity. Transient Ca²⁺ fluxes, mediated by Ca²⁺ permeable channels and decoded by Ca²⁺ binding sensor proteins, regulate various downstream cellular processes that are essential for both NLR-and PRR-mediated immunity (Wang and Luan, 2024; Weralupitiya *et al*., 2024). The late and long-lasting Ca²⁺ signature is quite singular for a biotic elicitor since, for the majority of MAMPs (Microbe-Associated Molecular Patterns) characterized to date, it has been shown that the Ca²⁺ influx is an early process (during the first few minutes) in plant immunity (Ranf *et al*., 2011). Interestingly, such a late and long signature is more reminiscent to ETI or some abiotic stresses (White and Broadley, 2003).

We found that RLs induce the expression of *AtCPK5* and *AtCPK6* genes. These CPKs are already known to be involved in PTI by regulating the expression of defense genes, most commonly through transcription factors of the WRKY family (Boudsocq *et al*., 2010; Zhou *et al*., 2020*b*, 2024). *At*CPK5 in particular is known to induce resistance to the pathogenic bacterium *Pst* DC3000 and the fungus *B. cinerea* (Sun *et al*., 2025).

RLs also activate an atypical ROS production signature in plants (Cloutier *et al*., 2021; Schellenberger *et al*., 2021). When ROS production was monitored in the three CPK-related mutants (*cpk5, cpk6 and cp5/6*) and compared to the parental line WT, it appeared that both *At*CPK5 and *At*CPK6 are involved in the regulation albeit to different degrees. Under our experimental conditions, ROS production in the *cpk5* background was slightly higher than in WT, suggesting that *At*CPK5 could act as a negative regulator of RL-induced ROS production. In the *cpk6* background, however, production did not differ from WT, providing no evidence for *At*CPK6 involvement in this process. However, in the *cpk5/6* mutant, ROS production was significantly higher than in both WT and *cpk6* suggesting that both *At*CPK5 and *At*CPK6 could act in synergy. *At*CPK5 and *At*CPK6 have previously been shown to act together. For instance, they function as positive, redundant regulators of *B. cinerea*-induced camalexin biosynthesis (Zhou *et al*., 2020*a*). Previous studies have proposed that *At*CPK5 and *At*CPK6 act positively to regulate the MAMP-induced ROS burst, particularly in response to elicitors such as flagellin 22 (flg22) (Boudsocq *et al*., 2010; Dubiella *et al*., 2013). A positive regulation by *At*CPK5/6 homologs is also found in other plant species. For example, *Os*CPK5/*Os*CPK13 and *St*CPK4/*St*CPK5 act as positive regulators of ROS production via *Os*RBOHD/*St*RBOHB phosphorylation in *Oryza sativa* and *Solanum tuberosum*, respectively (Kobayashi *et al*., 2007; Li *et al*., 2025). Our results also indicate that *At*CPK5 and *At*CPK6 could act in a complementary manner, but potentially as negative regulators in response to RLs. Interestingly, other CPKs, such as *At*CPK8 in Arabidopsis, have also been shown to negatively regulate oxidative stress. The *cpk8* mutant accumulated more ROS in leaves and stomatal guard cells compared with WT plants when treated with abscisic acid (ABA) (Zou *et al*., 2015). *At*CPK28 was also described as a negative regulator of MAMP-induced oxidative burst, increasing the turnover of Botrytis-Induced Kinase1 (BIK1 ; Monaghan *et al*., 2014), an enzyme that activates *At*RBOHD.

We also observed that RL-triggered activation of early and late defense genes was affected by *AtCPK5* and *AtCPK6* inactivation. Indeed, *AtWRKY46* expression was significantly higher in *cpk5/6* mutants. The three treated mutants showed similar levels of *AtFRK1* expression and *AtPR1* expression levels are highest in *cpk5* and *cpk5/6* mutants. This supports the hypothesis that *At*CPK5 and *At*CPK6 may function as negative regulators of defense gene expression in response to RLs in a redundant way, and both must be absent for an effect to be seen. They have previously been identified as partially redundant positive regulators of defense responses activated by different MAMPs and Damage-Associated Molecular Patterns, such as flg22, elongation factor 18 (elf18), plant elicitor peptides 3 (PEP3), and oligogalacturonides (OGs) (Boudsocq *et al.,* 2010; Dubiella *et al*., 2013; Gravino *et al*., 2015). These two kinases have also various WRKY transcription factors as common phosphorylation substrates (Gao *et al*., 2013, 2014; Cui *et al*., 2020). A study showed that *At*CPK5/6 (mainly AtCPK5) influence the plant’s defense response triggered by 2-deoxyglucose (2DG), a glucose analog that can act as a stress signal (Yamada and Mine, 2024). The 2DG-induction of defense-related genes is differentially altered in *cpk5/6* showing decrease of *AtNHL10* (NDR1/HIN1-Like 10) and *AtPAD3* (Phytoalexin Deficient 3) induction but increase of the salicylic acid biosynthesis genes *AtPBS3* (AvrPphB Susceptible 3) and *AtSID2* (Salicylic acid Induction-Deficient 2), indicating the versatility of this type of kinases. *At*CPK5 and *At*CPK6 are typically described as positive regulators of immune responses. To date, these CPKs are not referenced in the literature as negative regulators of biotic stress in Arabidopsis, although, our data suggest that they may also act as negative modulators of specific defense gene expression in response to RLs. This context-dependent regulation underlines the complexity of CPK-mediated signaling. *AtWRKY46* and *AtPR1* genes are positively regulated by SA (Hu *et al*., 2012; Hussain *et al*., 2018). We observed that these two genes have similar expression profiles following RL treatment. Several reports have established that *At*CPK5 plays a pivotal role in modulating phytohormone biosynthesis, particularly the SA pathway (Dubiella *et al*., 2013; Li *et al*., 2018; Yang *et al*., 2020). Collectively, these findings suggest that *At*CPK5 could act as a component of Ca2+ signals to coordinate hormonal regulation of some defense gene expression in Arabidopsis following RL perception*. At*CPK6 is also referenced in the literature as a regulator of immunity in plants, in cooperation with *At*CPK5 (gene expression, ROS accumulation and ethylene biosynthesis ; Yip Delormel and Boudsocq, 2019). Indeed, under our experimental conditions, only the *AtFRK1* marker was affected in the *cpk6* mutant and *At*CPK6 appears to mainly play a compensatory/synergistical role with its homologue *At*CPK5 in RL-induced signaling.

The *cpk5/6* double mutant exhibits increased susceptibility to the pathogenic bacteria *Pst* DC3000, indicating that *At*CPK5 and *At*CPK6 act as positive regulators of basal resistance against this bacterial infection. Our findings are consistent with previous reports showing that these kinases contribute in duo to effective plant defense responses (Boudsocq *et al*., 2010). In the context of RL challenge, a protective effect against *Pst* DC3000 was clearly confirmed in WT plants, as previously demonstrated (Schellenberger *et al*., 2021). Interestingly, this resistance effect was not abolished in *cpk5*, *cpk6* or *cpk5/6* mutant lines. Therefore, although our data support the involvement of *At*CPK5 and *At*CPK6 in RL-triggered signaling events and some defense responses, the RL-induced local resistance is not affected. One explanation could be the involvement of other CPKs, or entirely distinct branches of the immune network. Direct resistance to a pathogen can differ from induced resistance triggered by specific elicitors. For example, RLs are known to activate both SA and JA/ethylene pathways in Arabidopsis, thereby triggering a resistance to necrotrophic and biotrophic pathogen agents (Sanchez *et al*., 2012). In this case, the classical dichotomy between pathogen with different lifestyles and phytohormones is not conserved.

Various studies have shown that RLs integrate into the plasma membrane in a manner reminiscent of mechanical stresses (Monnier *et al*., 2019; Crouzet *et al*., 2020). Consequently, it is plausible that the perception of RLs, despite their bacterial origin, is similar to an abiotic stress. Accordingly, RLs induce electrolyte leakage in Arabidopsis cells (fig. 6 and Schellenberger *et al*., 2021). It has been shown that WT plant roots induce a late peak (6 hours after treatment) of ROS after the induction of salt stress (Xie *et al*., 2011). This is consistent with the ROS production we observed after RL challenge. Several CPKs have previously been identified as negative regulators of abiotic stress responses. For instance, *At*CPK9 and *At*CPK33 negatively regulate ABA-induced stomatal closure by modulating anionic currents (Li *et al*., 2016; Chen *et al*., 2019), while *At*CPK21 has been described as a negative regulator of osmotic stress tolerance (Franz *et al*., 2011). This functional diversity among CPKs may help to explain why *At*CPK5 and *At*CPK6 could act differently in this context compared to their well-established roles in direct biotic stress responses.

In conclusion, in this study, we demonstrated that *At*CPK5 and *At*CPK6 can act as negative regulators of some RL-mediated immune responses including ROS production and some defense gene activation. Interestingly, despite these molecular changes, *AtCPK5* and *AtCPK6* inactivation did not change the resistance to a biotrophic pathogen, suggesting that other factors are involved in RL-mediated plant resistance. These findings highlight the complexity of *At*CPK5 and *At*CPK6 and more generally CPK networks in plant immunity and provide a basis for further studies aiming to identify complementary regulators involved in RL-triggered plant resistance.

## Author contributions

**Conceptualization:** Stéphan Dorey, Sylvain Cordelier, Olivier Fernandez, Marie Boudsocq

**Data curation:** Juliette Stanek, Sylvain Cordelier, Olivier Fernandez

**Formal analysis:** Juliette Stanek, Olivier Fernandez, Dina Aggad, Sandrine Dhondt Cordelier, Jérôme Crouzet

**Funding acquisition:** Stéphan Dorey, Sylvain Cordelier, Olivier Fernandez

**Investigation:** Juliette Stanek, Dina Aggad, Sandra Villaume, Laetitia Parent

**Methodology:** Juliette Stanek, Sandra Villaume, Dina Aggad, Sandrine Dhondt Cordelier, Jérôme Crouzet

**Project administration:** Stéphan Dorey, Sylvain Cordelier, Olivier Fernandez

**Resources:** Stéphan Dorey, Sylvain Cordelier, Olivier Fernandez

**Supervision:** Stéphan Dorey, Sylvain Cordelier, Olivier Fernandez

**Validation:** Stéphan Dorey, Sylvain Cordelier, Olivier Fernandez

**Visualization:** Juliette Stanek

**Writing - original draft:** Juliette Stanek, Sylvain Cordelier, Olivier Fernandez

**Writing - review & editing:** Juliette Stanek, Stéphan Dorey, Sylvain Cordelier, Olivier Fernandez, Marie Boudsocq, Sandrine Dhondt Cordelier, Jérôme Crouzet

## Supplemental Material

**Supplemental Table 1:**
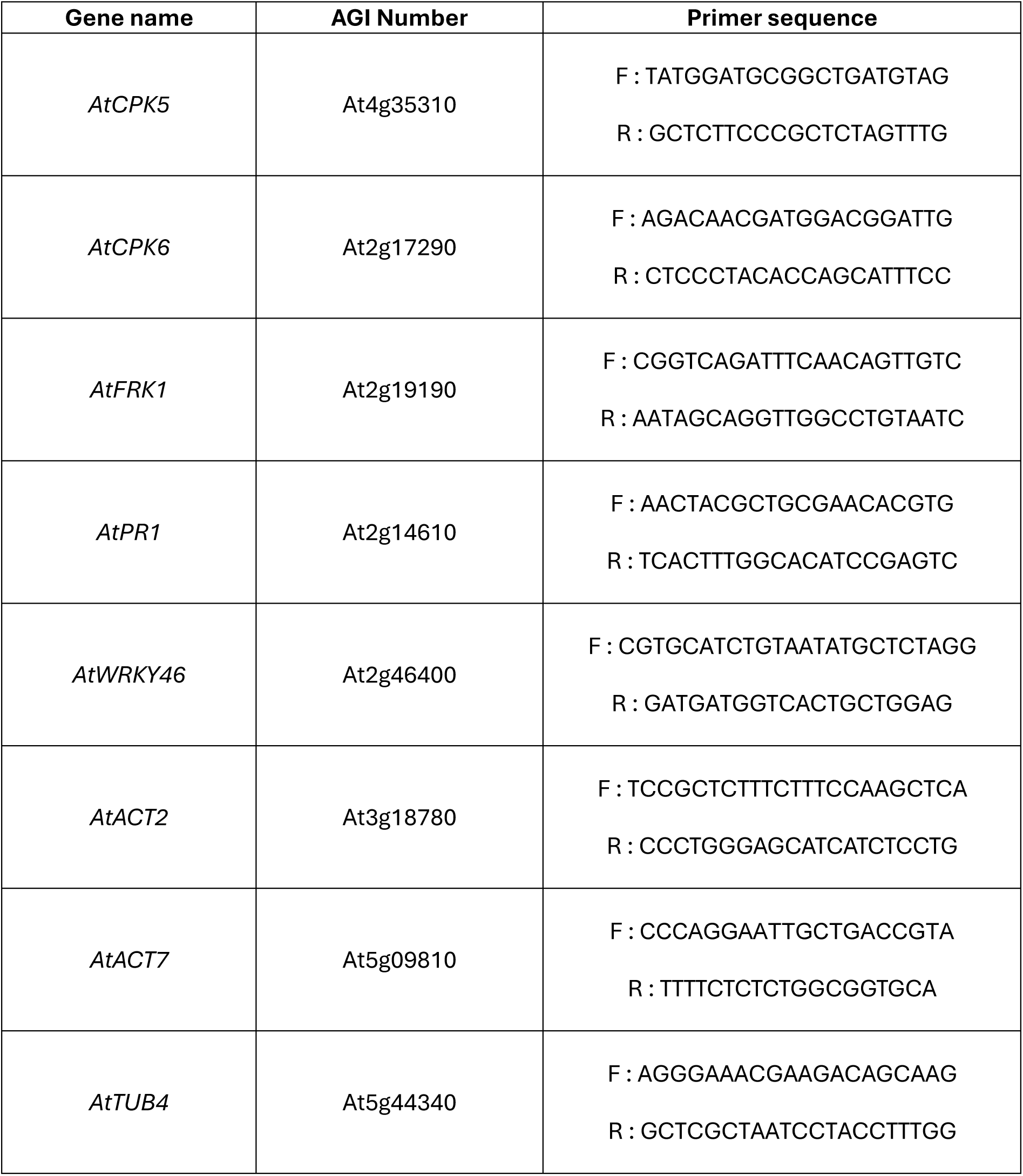
Sequences of primers used for qRT-PCR analyses. F : sequence of forward primer; R: sequence of reverse primer

## References

Aldon D, Mbengue M, Mazars C, Galaud J-P. 2018. Calcium Signalling in Plant Biotic Interactions. International Journal of Molecular Sciences 19, 665.

Biniaz Y, Tahmasebi A, Tahmasebi A, Albrectsen BR, Poczai P, Afsharifar A. 2022. Transcriptome Meta-Analysis Identifies Candidate Hub Genes and Pathways of Pathogen Stress Responses in Arabidopsis thaliana. Biology 11, 1155.

Blomberg J, Tasselius V, Vergara A, Karamat F, Imran QM, Strand Å, Rosvall M, Björklund S. 2024. Pseudomonas syringae infectivity correlates to altered transcript and metabolite levels of Arabidopsis mediator mutants. Scientific Reports 14, 6771.

Boudsocq M, Sheen J. 2010. Stress Signaling II: Calcium Sensing and Signaling. In: Pareek A, Sopory SK, Bohnert HJ, eds. Abiotic Stress Adaptation in Plants. Dordrecht: Springer Netherlands, 75–90.

Boudsocq M, Willmann MR, McCormack M, Lee H, Shan L, He P, Bush J, Cheng S-H, Sheen J. 2010. Differential innate immune signalling via Ca2+ sensor protein kinases. Nature 464, 418– 422.

Bredow M, Monaghan J. 2019. Regulation of Plant Immune Signaling by Calcium-Dependent Protein Kinases. Molecular Plant-Microbe Interactions® 32, 6–19.

Cao Y, Dai Y, Cui S, Ma L. 2008. Histone H2B Monoubiquitination in the Chromatin of FLOWERING LOCUS C Regulates Flowering Time in Arabidopsis. The Plant Cell 20, 2586–2602.

Chen D-H, Liu H-P, Li C-L. 2019. Calcium-dependent protein kinase CPK9 negatively functions in stomatal abscisic acid signaling by regulating ion channel activity in Arabidopsis. Plant Molecular Biology 99, 113–122.

Cloutier M, Prévost M-J, Lavoie S, et al. 2021. Total synthesis, isolation, surfactant properties, and biological evaluation of ananatosides and related macrodilactone-containing rhamnolipids. Chemical Science 12, 7533–7546.

Coca M, San Segundo B. 2010. AtCPK1 calcium-dependent protein kinase mediates pathogen resistance in Arabidopsis. The Plant Journal 63, 526–540.

Cordelier S, Crouzet J, Gilliard G, Dorey S, Deleu M, Dhondt-Cordelier S. 2022. Deciphering the role of plant plasma membrane lipids in response to invasion patterns: how could biology and biophysics help? (S Williams, Ed.). Journal of Experimental Botany 73, 2765–2784.

Crouzet J, Arguelles-Arias A, Dhondt-Cordelier S, et al. 2020. Biosurfactants in Plant Protection Against Diseases: Rhamnolipids and Lipopeptides Case Study. Frontiers in Bioengineering and Biotechnology 8.

Cui X, Zhao P, Liang W, et al. 2020. A Rapeseed WRKY Transcription Factor Phosphorylated by CPK Modulates Cell Death and Leaf Senescence by Regulating the Expression of ROS and SA-Synthesis-Related Genes. Journal of Agricultural and Food Chemistry 68, 7348–7359.

Dekomah SD, Bi Z, Dormatey R, Wang Y, Haider FU, Sun C, Yao P, Bai J. 2022. The role of CDPKs in plant development, nutrient and stress signaling. Frontiers in Genetics 13.

Du B, Haensch R, Alfarraj S, Rennenberg H. 2024. Strategies of plants to overcome abiotic and biotic stresses. Biological Reviews 99, 1524–1536.

Dubiella U, Seybold H, Durian G, Komander E, Lassig R, Witte C-P, Schulze WX, Romeis T. 2013. Calcium-dependent protein kinase/NADPH oxidase activation circuit is required for rapid defense signal propagation. Proceedings of the National Academy of Sciences 110, 8744–8749.

Franz S, Ehlert B, Liese A, Kurth J, Cazalé A-C, Romeis T. 2011. Calcium-Dependent Protein Kinase CPK21 Functions in Abiotic Stress Response in Arabidopsis thaliana. Molecular Plant 4, 83–96.

Furlan AL, Laurin Y, Botcazon C, Rodríguez-Moraga N, Rippa S, Deleu M, Lins L, Sarazin C, Buchoux S. 2020. Contributions and Limitations of Biophysical Approaches to Study of the Interactions between Amphiphilic Molecules and the Plant Plasma Membrane. Plants 9, 648.

Gao X, Chen X, Lin W, et al. 2013. Bifurcation of Arabidopsis NLR Immune Signaling via Ca2+-Dependent Protein Kinases. PLoS Pathogens 9, e1003127.

Gao X, Cox Jr. KL, He P. 2014. Functions of Calcium-Dependent Protein Kinases in Plant Innate Immunity. Plants 3, 160–176.

Gravino M, Savatin DV, Macone A, De Lorenzo G. 2015. Ethylene production in Botrytis cinerea- and oligogalacturonide-induced immunity requires calcium-dependent protein kinases. The Plant Journal 84, 1073–1086.

Hao X, Wang S, Fu Y, Liu Y, Shen H, Jiang L, McLamore ES, Shen Y. 2024. The WRKY46–MYC2 module plays a critical role in E-2-hexenal-induced anti-herbivore responses by promoting flavonoid accumulation. Plant Communications 5, 100734.

Harmon AC, Gribskov M, Gubrium E, Harper JF. 2001. The CDPK superfamily of protein kinases. New Phytologist 151, 175–183.

He P, Shan L, Lin N-C, Martin GB, Kemmerling B, Nürnberger T, Sheen J. 2006. Specific Bacterial Suppressors of MAMP Signaling Upstream of MAPKKK in Arabidopsis Innate Immunity. Cell 125, 563–575.

Hu Y, Dong Q, Yu D. 2012. Arabidopsis WRKY46 coordinates with WRKY70 and WRKY53 in basal resistance against pathogen Pseudomonas syringae. Plant Science 185–186, 288–297.

Hussain RMF, Sheikh AH, Haider I, Quareshy M, Linthorst HJM. 2018. Arabidopsis WRKY50 and TGA Transcription Factors Synergistically Activate Expression of PR1. Frontiers in Plant Science 9.

Jones JDG, Staskawicz BJ, Dangl JL. 2024. The plant immune system: From discovery to deployment. Cell 187, 2095–2116.

Kadota Y, Sklenar J, Derbyshire P, et al. 2014. Direct Regulation of the NADPH Oxidase RBOHD by the PRR-Associated Kinase BIK1 during Plant Immunity. Molecular Cell 54, 43–55.

Kiselev KV, Dubrovina AS. 2025. The role of calcium-dependent protein kinase (CDPK) genes in plant stress resistance and secondary metabolism regulation. Plant Growth Regulation doi: 10.1007/s10725-025-01295-6.

Kobayashi M, Ohura I, Kawakita K, Yokota N, Fujiwara M, Shimamoto K, Doke N, Yoshioka H. 2007. Calcium-Dependent Protein Kinases Regulate the Production of Reactive Oxygen Species by Potato NADPH Oxidase. The Plant Cell 19, 1065–1080.

Körber N, Bus A, Li J, Parkin IAP, Wittkop B, Snowdon RJ, Stich B. 2016. Agronomic and Seed Quality Traits Dissected by Genome-Wide Association Mapping in Brassica napus. Frontiers in Plant Science 7.

Kovács S, Nagy Á, Major G, Flors V, Rácz A, Mauch-Mani B, Jakab G. 2025. The role of PRLIP2 in the defence and growth of Arabidopsis Insertional mutagenesis of PRLIP2 gene alters hormone balance and defence responses in Arabidopsis. Journal of Plant Physiology 314, 154609.

Kudla J, Becker D, Grill E, Hedrich R, Hippler M, Kummer U, Parniske M, Romeis T, Schumacher K. 2018. Advances and current challenges in calcium signaling. New Phytologist 218, 414–431.

Kutschera A, Dawid C, Gisch N, et al. 2019. Bacterial medium-chain 3-hydroxy fatty acid metabolites trigger immunity in Arabidopsis plants. Science 364, 178–181.

Laird J, Armengaud P, Giuntini P, Laval V, Milner JJ. 2004. Inappropriate annotation of a key defence marker in Arabidopsis: will the real PR-1 please stand up? Planta 219, 1089–1092.

Li S, Han X, Yang L, Deng X, Wu H, Zhang M, Liu Y, Zhang S, Xu J. 2018. Mitogen-activated protein kinases and calcium-dependent protein kinases are involved in wounding-induced ethylene biosynthesis in. Plant, Cell & Environment 41, 134–147.

Li J, Ishii T, Yoshioka M, Hino Y, Nomoto M, Tada Y, Yoshioka H, Takahashi H, Yamauchi T, Nakazono M. 2025. CDPK5 and CDPK13 play key roles in acclimation to low oxygen through the control of RBOH-mediated ROS production in rice. Plant Physiology 197, kiae293.

Li C-L, Wang M, Wu X-M, Chen D-H, Lv H-J, Shen J-L, Qiao Z, Zhang W. 2016. THI1, a Thiamine Thiazole Synthase, Interacts with Ca2+-Dependent Protein Kinase CPK33 and Modulates the S-Type Anion Channels and Stomatal Closure in Arabidopsis. Plant Physiology 170, 1090–1104.

Liese A, Eichstädt B, Lederer S, Schulz P, Oehlschläger J, Matschi S, Feijó JA, Schulze WX, Konrad KR, Romeis T. 2024. Imaging of plant calcium-sensor kinase conformation monitors real time calcium-dependent decoding *in planta*. The Plant Cell 36, 276–297.

Liese A, Romeis T. 2013. Biochemical regulation of *in vivo* function of plant calcium-dependent protein kinases (CDPK). Biochimica et Biophysica Acta (BBA) - Molecular Cell Research 1833, 1582–1589.

Liu Y, Zhang H, Wang J, et al. 2024. Nonpathogenic Pseudomonas syringae derivatives and its metabolites trigger the plant “cry for help” response to assemble disease suppressing and growth promoting rhizomicrobiome. Nature Communications 15, 1907.

Mariyam S, Bhardwaj R, Khan NA, Sahi SV, Seth CS. 2023. Review on nitric oxide at the forefront of rapid systemic signaling in mitigation of salinity stress in plants: Crosstalk with calcium and hydrogen peroxide. Plant Science 336, 111835.

Monaghan J, Matschi S, Shorinola O, et al. 2014. The Calcium-Dependent Protein Kinase CPK28 Buffers Plant Immunity and Regulates BIK1 Turnover. Cell Host & Microbe 16, 605–615.

Monnier N, Furlan A, Botcazon C, Dahi A, Mongelard G, Cordelier S, Clément C, Dorey S, Sarazin C, Rippa S. 2018. Rhamnolipids From Pseudomonas aeruginosa Are Elicitors Triggering Brassica napus Protection Against Botrytis cinerea Without Physiological Disorders. Frontiers in Plant Science 9.

Monnier N, Furlan AL, Buchoux S, Deleu M, Dauchez M, Rippa S, Sarazin C. 2019. Exploring the Dual Interaction of Natural Rhamnolipids with Plant and Fungal Biomimetic Plasma Membranes through Biophysical Studies. International Journal of Molecular Sciences 20, 1009.

Mu Z, Xu M, Manda T, Yang L, Hwarari D, Zhu F-Y. 2024. Genomic survey and evolution analysis of calcium-dependent protein kinases in plants and their stress-responsive patterns in populus. BMC Genomics 25, 1108.

Ngou BPM, Ding P, Jones JDG. 2022a. Thirty years of resistance: Zig-zag through the plant immune system. The Plant Cell 34, 1447–1478.

Ngou BPM, Jones JDG, Ding P. 2022b. Plant immune networks. Trends in Plant Science 27, 255– 273.

Ngou BPM, Wyler M, Schmid MW, Kadota Y, Shirasu K. 2024. Evolutionary trajectory of pattern recognition receptors in plants. Nature Communications 15, 308.

Pečenková T, Pejchar P, Moravec T, Drs M, Haluška S, Šantrůček J, Potocká A, Žárský V, Potocký M. 2022. Immunity functions of Arabidopsis pathogenesis-related 1 are coupled but not confined to its C-terminus processing and trafficking. Molecular Plant Pathology 23, 664–678.

Ranf S, Eschen-Lippold L, Pecher P, Lee J, Scheel D. 2011. Interplay between calcium signalling and early signalling elements during defence responses to microbe- or damage-associated molecular patterns. The Plant Journal 68, 100–113.

Ranf S, Gisch N, Schäffer M, et al. 2015. A lectin S-domain receptor kinase mediates lipopolysaccharide sensing in Arabidopsis thaliana. Nature Immunology 16, 426–433.

Ravi B, Foyer CH, Pandey GK. 2023. The integration of reactive oxygen species (ROS) and calcium signalling in abiotic stress responses. doi: 10.1111/pce.14596.

Sanchez L, Courteaux B, Hubert J, Kauffmann S, Renault J-H, Clément C, Baillieul F, Dorey S. 2012. Rhamnolipids Elicit Defense Responses and Induce Disease Resistance against Biotrophic, Hemibiotrophic, and Necrotrophic Pathogens That Require Different Signaling Pathways in Arabidopsis and Highlight a Central Role for Salicylic Acid. Plant Physiology 160, 1630–1641.

Schellenberger R, Crouzet J, Nickzad A, et al. 2021. Bacterial rhamnolipids and their 3-hydroxyalkanoate precursors activate *Arabidopsis* innate immunity through two independent mechanisms. Proceedings of the National Academy of Sciences 118, e2101366118.

Schellenberger R, Touchard M, Clément C, Baillieul F, Cordelier S, Crouzet J, Dorey S. 2019. Apoplastic invasion patterns triggering plant immunity: plasma membrane sensing at the frontline. Molecular Plant Pathology 20, 1602–1616.

Smith JM, Heese A. 2014. Rapid bioassay to measure early reactive oxygen species production in Arabidopsis leave tissue in response to living Pseudomonas syringae. Plant Methods 10, 6.

Sun C, Chen Y, Ma A, et al. 2025. The kinase CPK5 phosphorylates MICRORCHIDIA1 to promote broad-spectrum disease resistance. The Plant Cell 37, koaf051.

Varnier A-L, Sanchez L, Vatsa P, et al. 2009. Bacterial rhamnolipids are novel MAMPs conferring resistance to Botrytis cinerea in grapevine. Plant, Cell & Environment 32, 178–193.

Verma S, Negi NP, Narwal P, Kumari P, Kisku AV, Gahlot P, Mittal N, Kumar D. 2022. Calcium signaling in coordinating plant development, circadian oscillations and environmental stress responses in plants. Environmental and Experimental Botany 201, 104935.

Wang R, He F, Ning Y, Wang G-L. 2020. Fine-Tuning of RBOH-Mediated ROS Signaling in Plant Immunity. Trends in Plant Science 25, 1060–1062.

Wang C, Luan S. 2024. Calcium homeostasis and signaling in plant immunity. Current Opinion in Plant Biology 77, 102485.

Wang J-P, Xu Y-P, Munyampundu J-P, Liu T-Y, Cai X-Z. 2016. Calcium-dependent protein kinase (CDPK) and CDPK-related kinase (CRK) gene families in tomato: genome-wide identification and functional analyses in disease resistance. Molecular Genetics and Genomics 291, 661–676.

Wdowiak A, Podgórska A, Szal B. 2024. Calcium in plants: an important element of cell physiology and structure, signaling, and stress responses. Acta Physiologiae Plantarum 46, 108.

Weralupitiya C, Eccersall S, Meisrimler C-N. 2024. Shared signals, different fates: Calcium and ROS in plant PRR and NLR immunity. Cell Reports 43.

White PJ, Broadley MR. 2003. Calcium in Plants. Annals of Botany 92, 487–511.

Xie Y-J, Xu S, Han B, Wu M-Z, Yuan X-X, Han Y, Gu Q, Xu D-K, Yang Q, Shen W-B. 2011. Evidence of Arabidopsis salt acclimation induced by up-regulation of HY1 and the regulatory role of RbohD-derived reactive oxygen species synthesis. The Plant Journal 66, 280–292.

Xu H, Liang X, Lloyd JR, Chen Y. 2024. Visualizing calcium-dependent signaling networks in plants. Trends in Plant Science 29, 117–119.

Yamada K, Mine A. 2024. Sugar coordinates plant defense signaling. Science Advances 10, eadk4131.

Yang L, Zhang Y, Guan R, Li S, Xu X, Zhang S, Xu J. 2020. Co-regulation of indole glucosinolates and camalexin biosynthesis by CPK5/CPK6 and MPK3/MPK6 signaling pathways. Journal of Integrative Plant Biology 62, 1780–1796.

Yang P, Zhao L, Gao YG, Xia Y. 2023. Detection, Diagnosis, and Preventive Management of the Bacterial Plant Pathogen Pseudomonas syringae. Plants 12, 1765.

Yip Delormel T, Boudsocq M. 2019. Properties and functions of calcium-dependent protein kinases and their relatives in Arabidopsis thaliana. New Phytologist 224, 585–604.

Yuan S, Jiang H, Wang Y, Zhang L, Shi Z, Jiao L, Meng D. 2025. A 3R-MYB transcription factor is involved in Methyl Jasmonate-Induced disease resistance in Agaricus bisporus and has implications for disease resistance in Arabidopsis. Journal of Advanced Research 73, 117–131.

Zhang H, Zhu J, Gong Z, Zhu J-K. 2022. Abiotic stress responses in plants. Nature Reviews Genetics 23, 104–119.

Zhou X, Lei Z, An P. 2024. Post-Translational Modification of WRKY Transcription Factors. Plants 13, 2040.

Zhou J, Wang X, He Y, Sang T, Wang P, Dai S, Zhang S, Meng X. 2020a. Differential Phosphorylation of the Transcription Factor WRKY33 by the Protein Kinases CPK5/CPK6 and MPK3/MPK6 Cooperatively Regulates Camalexin Biosynthesis in Arabidopsis. The Plant Cell 32, 2621–2638.

Zhou J, Wang X, He Y, Sang T, Wang P, Dai S, Zhang S, Meng X. 2020b. Differential Phosphorylation of the Transcription Factor WRKY33 by the Protein Kinases CPK5/CPK6 and MPK3/MPK6 Cooperatively Regulates Camalexin Biosynthesis in Arabidopsis. The Plant Cell 32, 2621–2638.

Zia R, Nawaz MS, Siddique MJ, Hakim S, Imran A. 2021. Plant survival under drought stress: Implications, adaptive responses, and integrated rhizosphere management strategy for stress mitigation. Microbiological Research 242, 126626.

Zou J-J, Li X-D, Ratnasekera D, Wang C, Liu W-X, Song L-F, Zhang W-Z, Wu W-H. 2015. Arabidopsis CALCIUM-DEPENDENT PROTEIN KINASE8 and CATALASE3 Function in Abscisic Acid-Mediated Signaling and H2O2 Homeostasis in Stomatal Guard Cells under Drought Stress. The Plant Cell 27, 1445–1460.

